# Regulation of midgut cell proliferation impacts *Aedes aegypti* susceptibility to dengue virus

**DOI:** 10.1101/271288

**Authors:** Mabel L. Taracena, Vanessa Bottino-Rojas, Octavio A.C. Talyuli, Ana Beatriz Walter-Nuno, José Henrique M. Oliveira, Yesseinia I. Angleró-Rodriguez, Michael B. Wells, George Dimopoulos, Pedro L. Oliveira, Gabriela O. Paiva-Silva

**Affiliations:** Laboratório de Bioquímica de Artrópodes Hematófagos, Instituto de Bioquímica Médica Leopoldo de Meis, Programa de Biologia Molecular e Biotecnologia, Universidade Federal do Rio de Janeiro, Rio de Janeiro, Brasil, 21941-902; Instituto Nacional de Ciência e Tecnologia em Entomologia Molecular (INCT-EM), Brasil; Departamento de Microbiologia, Imunologia e Parasitologia. Universidade Federal de Santa Catarina. Florianópolis, SC, Brasil, 88040-970.; W. Harry Feinstone Department of Molecular Microbiology and Immunology, Bloomberg School of Public Health, Johns Hopkins University, Baltimore, MD 21205, USA; Department of Cell Biology, Johns Hopkins University School of Medicine, Baltimore, MD 21205, USA; Johns Hopkins Malaria Research Institute, Johns Hopkins Bloomberg School of Public Health, Baltimore, MD 21205, USA.

**Keywords:** *Aedes aegypti*, Intestinal Stem Cell, Dengue virus, viral infection, midgut homeostasis, strain susceptibility, RNAi, Notch ligand Delta, vector competence

## Abstract

*Aedes aegypti* is the vector of some of the most important vector-borne diseases like Dengue, Chikungunya, Zika and Yellow fever, affecting millions of people worldwide. The cellular processes that follow a blood meal in the mosquito midgut are directly associated with pathogen transmission. We studied the homeostatic response of the midgut against oxidative stress, as well as bacterial and dengue virus (DENV) infections, focusing on the proliferative ability of the intestinal stem cells (ISC). Inhibition of the peritrophic matrix (PM) formation led to an increase in ROS production by the epithelial cells in response to contact with the resident microbiota, suggesting that maintenance of low levels of ROS in the intestinal lumen is key to keep ISCs division in balance. We show that dengue virus infection induces midgut cell division in both DENV susceptible (Rockefeller) and refractory (Orlando) mosquito strains. However, the susceptible strain delays the activation of the regeneration process compared with the refractory strain. Impairment of the Delta/Notch signaling, by silencing the Notch ligand Delta using RNAi, significantly increased the susceptibility of the refractory strains to DENV infection of the midgut. We propose that this cell replenishment is essential to control viral infection in the mosquito. Our study demonstrates that the intestinal epithelium of the blood fed mosquito is able to respond and defend against different challenges, including virus infection. In addition, we provide unprecedented evidence that the activation of a cellular regenerative program in the midgut is important for the determination of the mosquito vectorial competence.

## Authors’ summary

*Aedes* mosquitoes are important vectors of arboviruses, representing a major threat to public health. While feeding on blood, mosquitoes address the challenges of digestion and preservation of midgut homeostasis. Damaged or senescent cells must be constantly replaced by new cells to maintain midgut epithelial integrity. In this study, we show that the intestinal stem cells (ISCs) of blood-fed mosquitoes are able to respond to abiotic and biotic challenges. Exposing midgut cells to different types of stress, such as the inhibition of the peritrophic matrix formation, changes in the midgut redox state, or infection with entomopathogenic bacteria or viruses, resulted in an increased number of mitotic cells in blood-fed mosquitoes. Mosquito strains with different susceptibilities to DENV infection presented different time course of cell regeneration in response to viral infection. Knockdown of the Notch pathway in a refractory mosquito strain limited cell division after infection with DENV and resulted in increased mosquito susceptibility to the virus. Conversely, inducing midgut cell proliferation made a susceptible strain more resistant to viral infection. Therefore, the effectiveness of midgut cellular renewal during viral infection proved to be an important factor in vector competence. These findings can contribute to the understanding of virus-host interactions and help to develop more successful strategies of vector control.

## Introduction

The mosquito *Aedes aegypti* is a vector of several human pathogens, such as flaviviruses, including Yellow fever (YFV), Dengue (DENV) and Zika (ZIKV), and thus this mosquito exerts an enormous public health burden worldwide [1,2]. During the transmission cycle, these insects feed on volumes of blood that are 2-3 times their weight, and the digestion of this large meal results in several potentially damaging conditions [3]. The digestion of blood meal requires intense proteolytic activity in the midgut and results in the formation of potentially toxic concentrations of heme, iron, amino acids and ammonia [4]. The midgut is also the first site of interaction with potential pathogens, including viruses, and supports a dramatic increase in intestinal microbiota after blood feeding [5,6]. To overcome these challenges, the ingestion of a blood meal is followed by several physiological processes, such as formation of a peritrophic matrix (PM) [7,8] and down-regulation of reactive oxygen species (ROS) production. In addition, the midgut epithelium is the first barrier that viruses must cross in the mosquito to achieve a successful viral cycle (reviewed in [9]). Thus, in order to ensure epithelial integrity and the maintenance of midgut homeostasis, the midgut epithelium must fine tune key cellular mechanisms, including cell proliferation and differentiation.

In both vertebrate and invertebrate animals, the gut epithelia have a similar basic cellular composition: absorptive enterocytes (ECs) that represent the majority of the differentiated cells and are interspersed with hormone-producing enteroendocrine cells (ee). The intestinal stem cells (ISCs) and enteroblasts (EB) account for the progenitor cells, responsible for replenishing the differentiated cells that are lost due to damage or aging [10–14]. In *A. aegypti*, description of the different cellular types and functions started with identification and basic characterization of absorptive (ECs) and non-absorptive cells (ISC, EB, and enteroendocrine cells) [15]. To date, the study of division properties of the ISCs in this vector species remains limited to the description of the division process during metamorphosis [16].

Several conserved signaling pathways are known to be involved in midgut tissue renewal and differentiation. Comparative genomic analysis of some of these pathways has been done between *Drosophila melanogaster* and vector mosquitoes [17,18], but functional studies in *Aedes*, under the context of tissue regeneration, are still necessary. Notably, the Notch signaling pathway regulates cell differentiation in the midgut of both mammals and *Drosophila*. In *Drosophila*, loss of function of Notch is attributed to the increase of intestinal cell proliferation and tumor formation [19]. However, it has already been shown that depletion of Notch in *Drosophila* ISCs also leads to stem cell loss and premature ee cell formation [20]. Accordingly, disruption of Notch signaling in mice has resulted in decreased cell proliferation coupled with secretory cell hyperplasia, whereas hyperactivation of Notch signaling results in expanded proliferation with increased numbers of absorptive enterocytes [21], as also observed in *Drosophila* [20].

In *Drosophila*, the ingestion of cytotoxic agents, such as dextran sodium sulfate (DSS), bleomycin or paraquat, or infection by pathogenic bacteria can stimulates cell turnover, increasing the midgut ISC mitotic index [18,22]. Similar to that, it has been recently shown that cell damage produced by ingestion of several stressors also induced intestinal cell proliferation in sugar-fed *Aedes albopictus* [23]. Likewise, viral infections can trigger cellular responses, such as apoptosis or autophagy, in different infection models [24–27]. However, the interplay between intestinal cell proliferation and pathogen transmission has been a neglected subject in the literature.

In this study, we have characterized the dynamics of *A. aegypti* intestinal epithelium proliferation during blood meal digestion in response to oxidative stress, bacterial infections, and viral infections. We have also shown that two mosquito strains with different DENV susceptibilities [28] presented differences in cell mitotic rates after viral infection. Finally, our results indicate for the first time that the ability to replenish midgut cells by modulation of cell renewal involves the Delta-Notch signaling and is a key factor that influences *A. aegypti* competence to transmit DENV. We show that the cell proliferation rates influences mosquito infection and vector competence for DENV.

## Results

*Aedes aegypti* adult females acquire DENV and other arboviruses during the blood feedings that are needed to complete the reproductive cycle of the mosquito. To characterize the epithelial adaptation to this event, we first evaluated the cellular response to the blood meal itself. Upon ingestion, the blood induces dramatic changes in the Red strain mosquito midgut at a chemical, microbiological and physiological level. We attempted to dissect each of these challenges, to understand the delicate balance of the factors that play a role in the intestinal micro-environment in which the arbovirus has to thrive in order to pass to the salivary gland and be transmitted.

### Characterization of the adult intestinal cells and their regenerative capacity in *A. aegypti* adult midgut epithelium

The tissue homeostasis of the midgut depends on the ability to replenish the damaged cells, and this depends on the presence of ISCs. Due to the lack of specific markers for progenitor cells for *A. aegypti*, we used morphological and physiological parameters to define the presence of ISCs in the adult females. Progenitor cells are well characterized for their basal positioning and being diploid, different to the apical localization of differentiated cells and the polyploidy of enterocytes. Both cell types were clearly distinctive, as well as the peritrophic matrix, in the midgut epithelium of blood-fed adult females **(Fig 1A)**. The further characterization of ISC’s was performed with phospho-histone 3 antibodies, to specifically mark cells undergoing mitosis. In Fig 1B, it can be observed the two monolayers of the *A. aegypti* midgut, where ECs are clearly distinguishable and the PH3+ cell is found, with nuclei corresponding to the diploid size, located basally. Clearly, not every ISC present in the tissue is going to be found undergoing mitosis, but the presence of PH3+ cells, undoubtedly characterizes such cells as ISCs.

**Fig 1.**
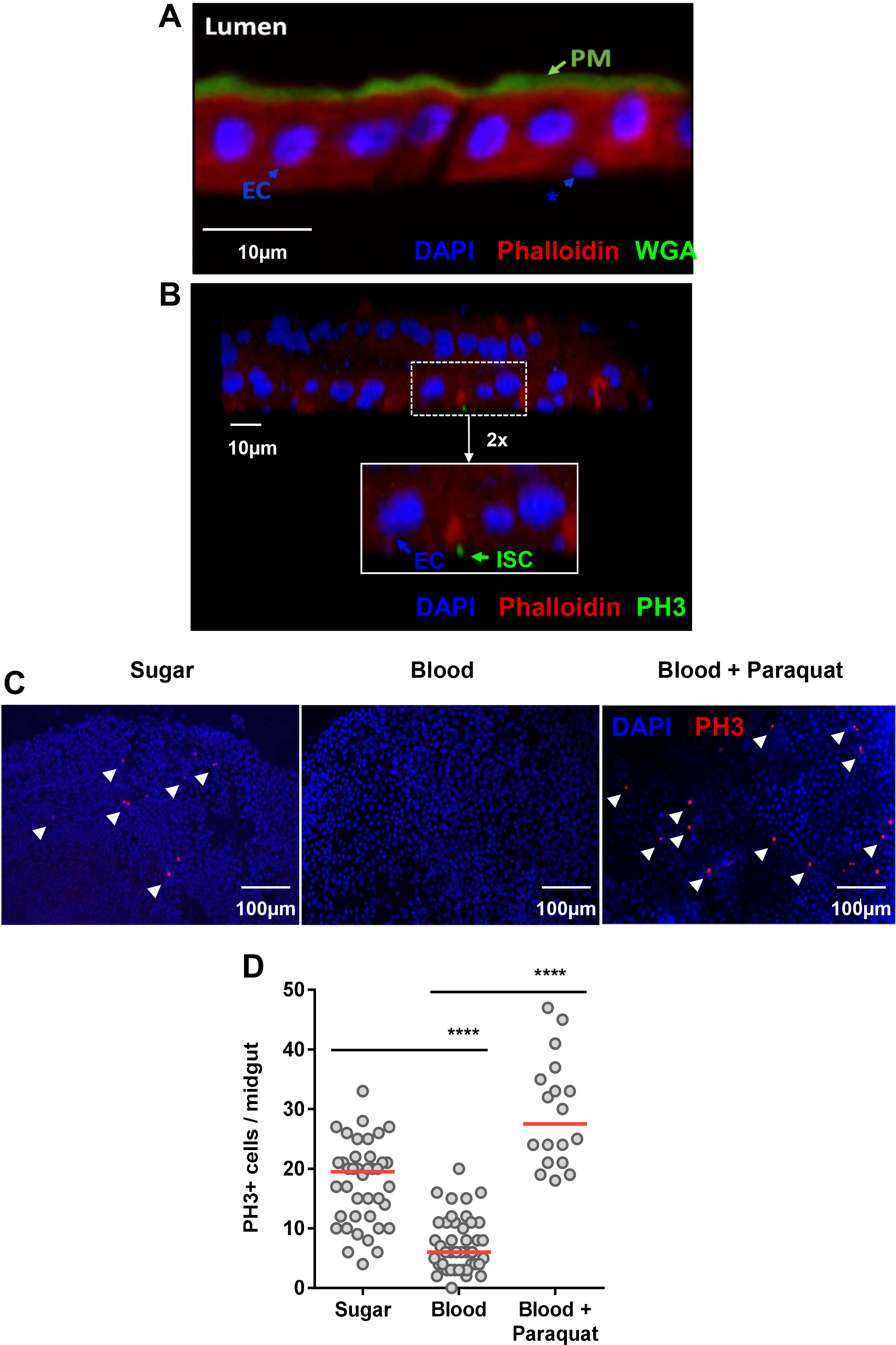
General structure of the midgut epithelium of *Aedes aegypti* and modulation of cell proliferation upon blood meal. The midgut epithelium from a blood-fed *A. aegypti* female was fixed in PFA, and in **(A)** sections of 0.14 μm were stained with WGA-FITC (green), red phalloidin (red) and DAPI (blue). The peritrophic matrix (PM), intestinal lumen (Lumen), polyploid enterocytes (EC) and basally localized – putative proliferative cells (*) – are visible. In **(B)**, confocal image (z-stack of 0.7 μm slides (20X)) of the two monolayers of the midgut of a blood-fed female, 5 days post feeding, stained with Ph3 mouse antibody (green), DAPI (blue), and phalloidin (red) – Inset (2x): polyploid enterocytes (EC) are Ph3-positive ISC (ISC) are visible. **(C)** Mosquitoes were fed on a sugar solution (10% sucrose), blood or blood supplemented with 100μM of the pro-oxidant paraquat. The insect midguts were dissected 24 hours after feeding and immunostained for PH3. Representative images of mitotic (PH3-labeled) cells (red) in the epithelial midgut of animals fed on sugar, blood or blood supplemented with paraquat are shown. The nuclei are stained with DAPI (blue). The arrowheads indicate PH3+ cells. **(D)** Quantification of PH3-positive cells per midgut of sugar, blood or blood plus paraquat-fed mosquitoes for sugar and blood and 18 for blood-paraquat fed midguts. The medians of at least three independent experiments are shown (n=40 for sugar and blood and n=18 for paraquat supplemented blood). The experiments were performed on Red Eye mosquito strain. The asterisks indicate significantly different values, **** P<0.0001 (Student’s t-test).

To evaluate the homeostatic cell proliferation of the *Aedes aegypti* midgut, we observed the number of cells undergoing mitosis in adult females. After a blood meal, the midgut epithelium showed a lower number of cells undergoing mitosis (phospho-histone 3 positive; PH3+) compared with that of sugar-fed insects **(Fig 1C and D)**. To test if this decrease in mitotic cells was due to progenitor cell impairment, we fed insects with blood supplemented with the pro-oxidant compound paraquat. The midgut epithelium responded to an oxidative challenge by increasing mitosis **(Fig 1C and D)**, indicating that the intestinal stem cells maintained the ability to divide and replenish damage cells after an insult at blood-fed conditions.

### Peritrophic matrix reduces cell proliferation induced by microbial infection

A hallmark of blood digestion is the formation of the peritrophic matrix (PM), a chitin and protein-rich non-cellular layer secreted by the midgut epithelium [7,8]. The mosquito PM surrounds the blood bolus, limiting a direct contact between the epithelium, the blood meal and the indigenous microbiota, thereby playing a similar function as the vertebrate digestive mucous layer. Ingestion of blood contaminated with bacteria allows close contact of these microorganisms to the midgut epithelium before PM formation, which is completed formed only a few hours (14 to 24 hours) after a blood meal [7]. In fact, oral infection with sub-lethal concentrations of the non-pathogenic *Serratia marcescens* or the entomopathogenic *Pseudomonas entomophila* bacteria resulted in a significant increase in mitosis of the epithelium cells **(Fig 2A and B)**. The increased cell turnover was also observed when heat-killed *P. entomophila* was provided through the blood, indicating that molecules derived from these entomopathogenic bacteria are sufficient to trigger the cell proliferation program, not necessarily requiring tissue infection **(Fig 2B)**. In this case, tissue damage may at least partially be attributed to the lack of cell membrane integrity promoted by Monalysin, a pore-forming protein produced by *P. entomophila* [29].

**Fig 2.**
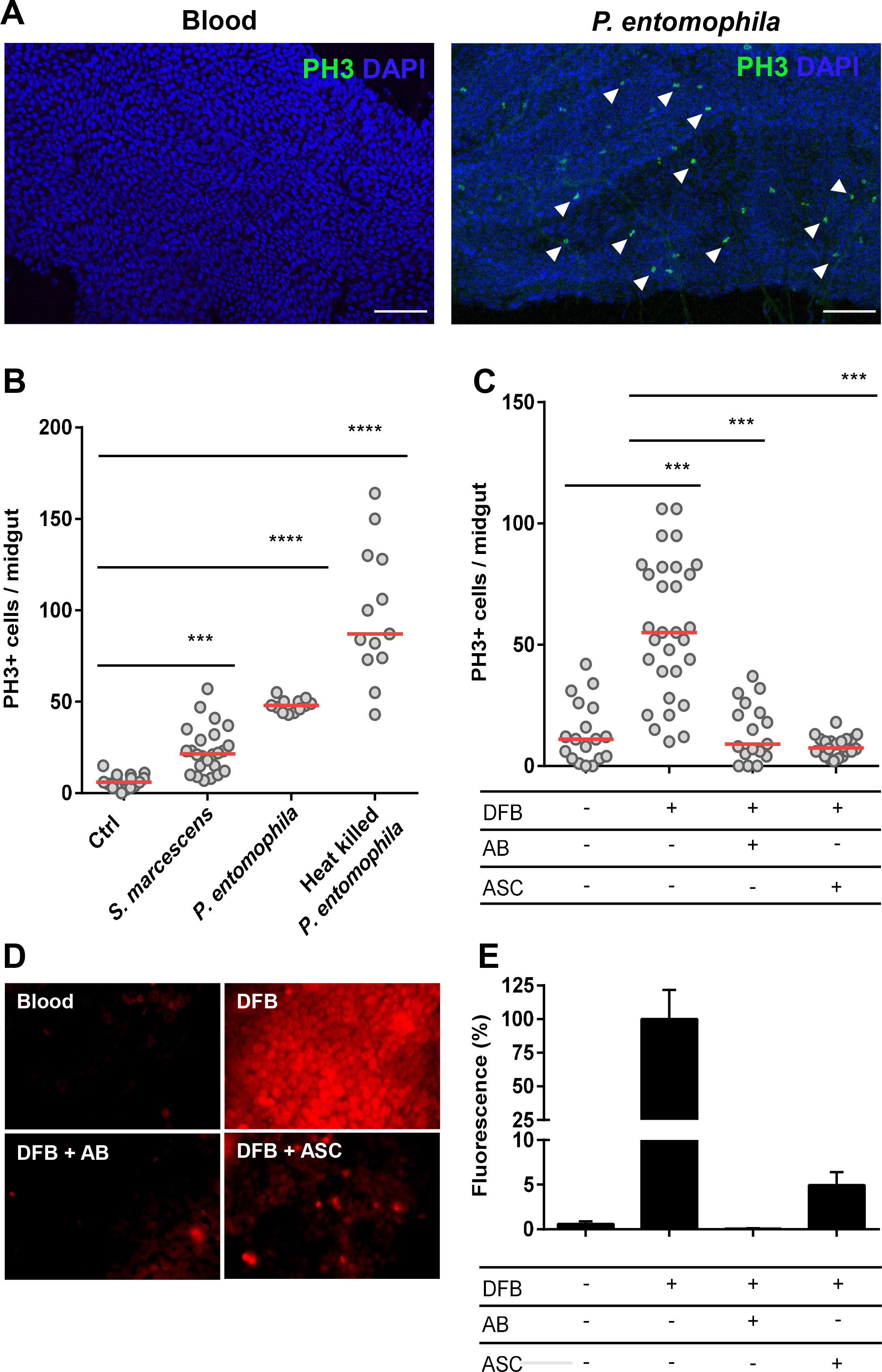
The peritrophic matrix shapes intestinal homeostasis by limiting contact of the gut epithelium with the microbiota and preventing ROS production. Red strain mosquitoes were fed on normal blood or blood infected with non-pathogenic *S. marcescens* or entomopathogenic *P. entomophila* bacteria. Another group of mosquitoes was fed blood supplemented with heat-killed *P. entomophila*. The midguts were dissected 24 hours after feeding and immunostained for PH3. **(A)** Representative images of PH3-labeled mitotic cells (green) of the midgut epithelium 24 h after a naïve blood meal or blood infected with *P. entomophila*. The nuclei are stained with DAPI (blue). The arrowheads indicate PH3+ cells. Scale bar=100 μm **(B)** Total PH3-positive cells were quantified from the midguts of mosquitoes fed on naïve and bacteria-infected blood (n=25) or heat-inactivated *P. entomophila*. (n=12). The medians of three independent experiments are shown. The asterisks indicate significantly different values *** P<0.001 and **** P<0.0001 (Student’s t-test). **(C)** Inhibition of PM formation results in a significant increase of progenitors cells under mitosis. The mosquitoes were fed blood or blood supplemented with diflubenzuron (DFB), DFB plus an antibiotic cocktail (AB) or DFB plus 50 mM ascorbate (ASC). The midguts were dissected 24 hours after feeding, and the mitotic indices were quantified by counting PH3+ cells. The medians of at least three independent experiments are shown (n=30). The asterisks indicate significantly different values *** P<0.001 and **** P<0.0001 (Student’s t-test). **(D)** Assessments of reactive oxygen species in the midguts were conducted by incubating midguts of insects fed as in **(C)** with a 50 μM concentration of the oxidant-sensitive fluorophore DHE. **(E)** Quantitative analysis of the fluorescence images shown in **(D)** were performed using ImageJ 1.45s software (n = 7 − 9 insects).

Supplementation of blood with diflubenzuron (DFB), a chitin synthesis inhibitor [30], leads to the inhibition of PM production, exposing the gut epithelium directly to the luminal content **(S1 Fig)**. Consequently, DFB administration resulted in elevated numbers of mitotic cells **(Fig 2C)**. The co-ingestion of antibiotics completely abolished this effect of DFB on cell proliferation (Fig 2C), demonstrating that in the absence of the microbiota, the lack of the peritrophic matrix did not result in elevated mitosis. These results indicate that not only oral infection with pathogenic bacteria, but also the proliferation of the resident microbiota (by inhibition of PM in this case), in contact with the epithelium, can trigger the midgut proliferative program.

Exposure of *Drosophila* enterocytes to bacteria results in ROS production as a microbiota control mechanism. However, the oxidative species produced as a result of bacterial presence can also cause damage to the midgut cells [31–34]. When mosquitoes were fed with blood supplemented with DFB together with the antioxidant ascorbate (ASC), the mitosis levels dropped significantly **(Fig 2C)**. The ROS production by the midgut epithelium was assessed by fluorescence microscopy using the fluorescent oxidant-sensing probe dihydroethidium (DHE). As shown in **Figures 2D and E**, the midguts of DFB-fed mosquitoes exhibited a high fluorescence signal, indicating an intense production of ROS. The intensity of the fluorescence signal of the DFB-treated midguts was significantly reduced upon ascorbate supplementation of the blood meal. Similarly, the suppression of microbiota with antibiotics dramatically reduced ROS levels. These results suggest a mechanism linking PM impairment to ISC proliferation, indicating that the direct exposure of the midgut epithelium to microbiota activates the production of ROS as part of an immune response.

### Infection with Dengue virus affects midgut epithelia regeneration

The role of epithelial tissue regeneration of the midgut upon viral infection has not been investigated in mosquitoes. Thus, we decided to evaluate the gut regeneration pattern of two mosquito strains that are known to exhibit different susceptibilities to DENV infection [28]. In basal conditions, i.e. sugar fed, all the strains used in this study presented no difference in the number of cells under mitosis **(S2 Fig)**. However, after 24 hours of taking a non-infected blood meal (day 1), the DENV refractory Orlando (Orl) strain presented a higher number of mitotic cells compared with the susceptible Rockefeller (Rock) strain **(Fig 3A and B)**, indicating that the refractory strain is naturally more proliferative than the susceptible one under these conditions. In the following days, both strains showed similar time course profiles of mitotic activity. Upon ingestion of DENV-infected blood, the refractory Orlando strain showed an increase of mitotic cells, peaking at the second day post blood meal **(Fig 3C)**. Subsequently, these midguts showed low numbers of cells in mitosis throughout the remaining course of infection, reaching a similar number as non-infected midguts. In contrast, the susceptible Rockefeller strain showed a delayed regenerative response, only reaching the maximum rate at five days after infection (Fig 3C). These results suggest that the midgut cells of refractory mosquitoes are able to respond more promptly to the early events of infection.

**Fig 3.**
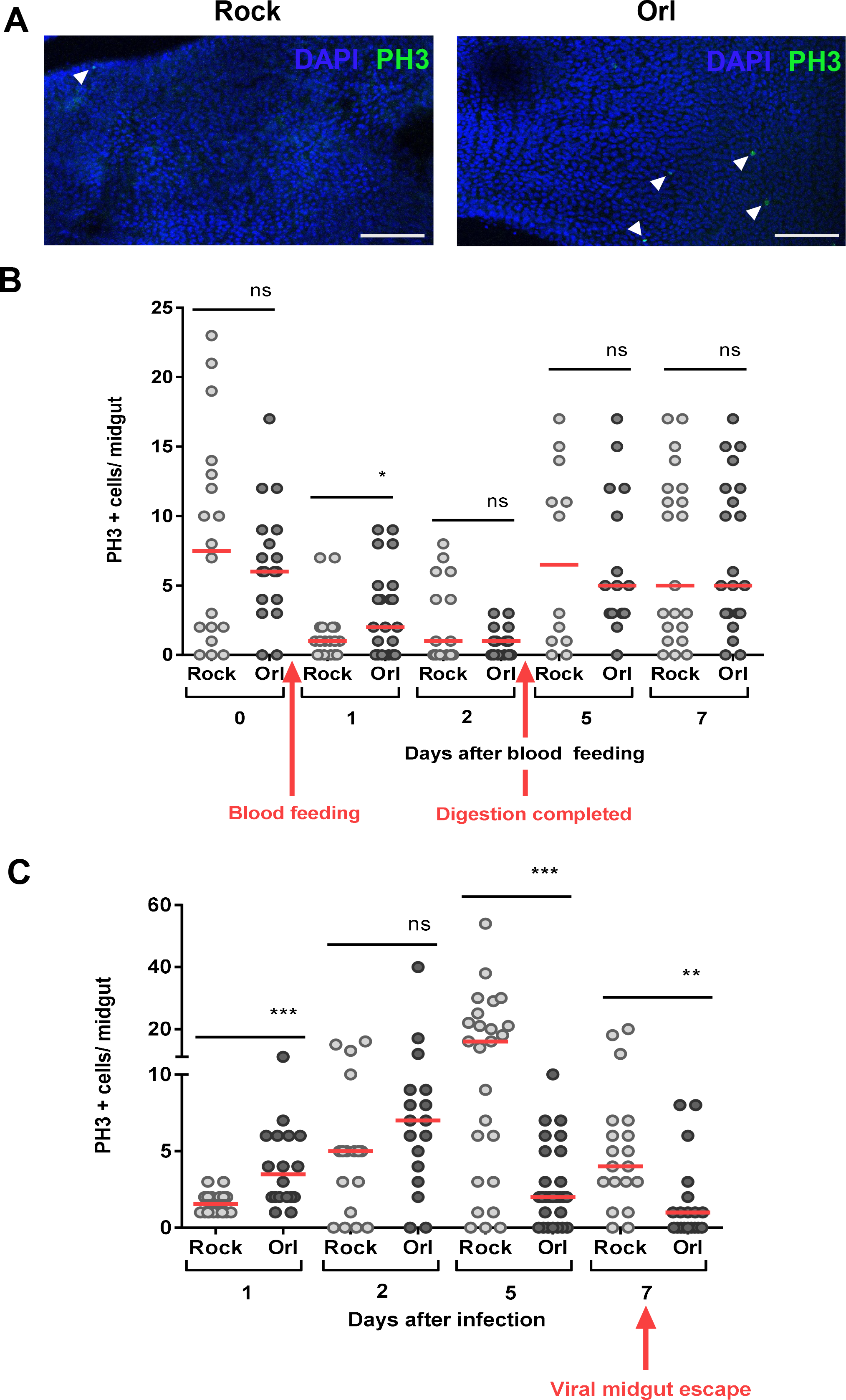
Dengue virus infection impacts midgut homeostasis in a strain specific manner. **(A)** Blood feeding induces different levels of PH3 positive cells in the midgut of the susceptible (Rock) and refractory (Orl) strains 24 hours after the meal. Representative images of PH3 labeling in both strains, 24 hours after the blood meal. The nuclei are stained with DAPI. The arrowheads indicate PH3+ cells. Scale bar=100 μm. **(B)** Mosquitoes from the two strains were blood fed and at day zero (non blood-fed) or at different days after feeding, the midguts were dissected and immunostained for PH3. The red arrows indicate the time of blood feeding and the time in which the digestion is completed (after blood bolus excretion).In **(C)** the mosquitoes were fed on DENV2-infected blood and mitotic-cell counting was performed at different days after infection. The red arrow indicates the time of DENV escape from the midgut to hemocoel. The medians of at least three independent experiments are shown (n=30). The asterisks indicate significantly different values * P<0.05 ** P<0.01 and *** P<0.001 (Student’s t-test).

To test whether the differences in gut homeostatic responses between the two strains could be a determinant of refractoriness/susceptibility, we disturbed the homeostatic condition of ISCs by silencing *delta* expression. The Notch ligand Delta (Dl) is an upstream component of the Notch pathway that is involved in cell division and differentiation. The *delta* gene is expressed in adult ISC cells. Thus, accumulation of Delta is used as a marker of ISCs in *Drosophila* [19]. Furthermore, Delta expression is induced by infection in the *Drosophila* midgut [35]. The efficiency and duration of Delta silencing by RNAi are shown in Figure 4A and Figure S3, respectively. Silencing *delta* led to a significant reduction in mitosis in both mosquito strains **(Fig 4B and C)**. Interestingly, silencing of *delta* did not have an effect on infection susceptibility in the Rockefeller strain **(Fig 4D)**. In contrast, it significantly increased susceptibility of the Orlando strain to DENV infection, as observed by the increased viral titers in the *delta*-silenced refractory strain compared with the dsGFP-injected group **(Fig 4D)**. Conversely, when the susceptible strain was pre-treated with DSS, a known inducer of midgut cell damage, and thereby ISC proliferation [18] and **Figure S4**, a significant reduction was seen in both DENV infection intensity **(Fig 4E)** and prevalence **(Fig 4F)** in the midgut, compared with non-treated mosquitoes. Similar results were observed when DSS-treated Rock mosquitoes were infected with DENV4 isolates **(Fig. S5)**. These data clearly indicate that the ability of midguts to respond at the cellular level, via regeneration of epithelial cells, modulates the success of viral infection of *A. aegypti*. Furthermore, these results show for the first time that the mosquito processes required to replenish damaged cells and control tissue homeostasis are determinants of vector competence.

**Fig 4.**
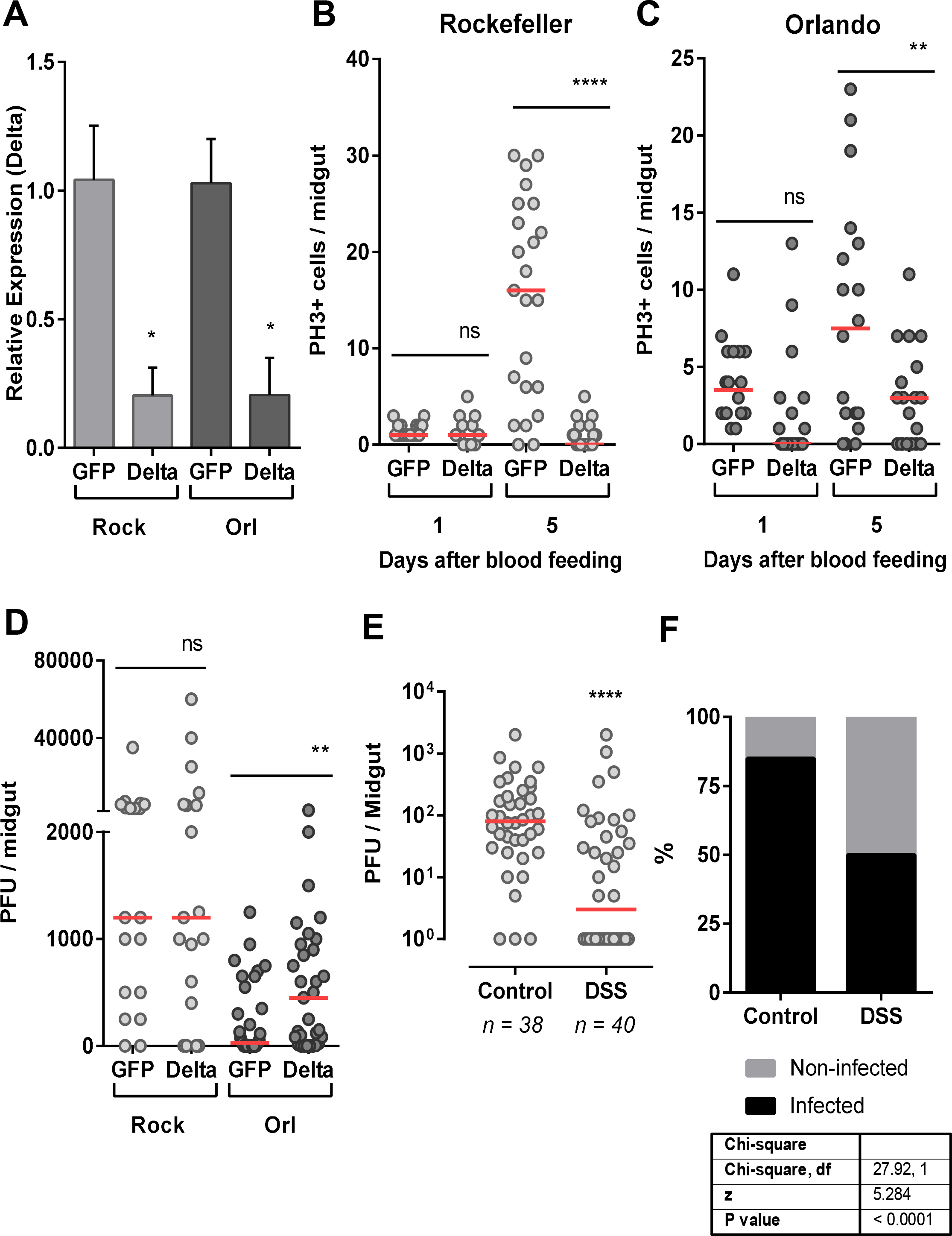
Interference in gut homeostatic response impacts vector competence. **(A)**. The midguts of dsRNA-injected Rockefeller and Orlando mosquitoes were dissected 24 days after a blood meal for silencing quantification of Delta, the ligand of Notch. Total PH3-positive cells were quantified from midguts of silenced Delta or control (GFP) mosquitoes from the Rockefeller **(B)** or Orlando **(C)** strains, both 1 and 5 days after blood meal. **(D)** dsRNA-Injected mosquitoes were fed DENV2-infected blood, and 5 days after the infection, the midguts were dissected for the plaque assay. **(E)** The susceptible (Rockefeller) mosquitoes were pre-treated with the tissue-damaging dextran sulfate sodium (DSS) accordingly to material and methods section. Twelve hours after the end of the DSS treatment, the mosquitoes were fed with DENV-2-infected blood. After 5 days, the midguts were dissected for the plaque assay. **(F)** The percentage of infected midguts (infection prevalence) was scored from the same set of data as in (E). The medians of at least three independent experiments are shown. n=20-25 in (A),(B) and (C);n=20-26 in (D) and n=40 in (E). Statistical analyzes used were: Student’s t-test for (A), (B) and (C); Mann-Whitney U-tests were used for infection intensity (D and E); and chi-square tests were performed to determine the significance of infection prevalence analysis (F). *P<0.05, ** P<0.01, **** P<0.0001.

## Discussion

Cell renewal is known to be the basis of midgut epithelial integrity in model animals such as fly and mice. Given the importance of the midgut epithelium in mosquitoes, where this tissue is effectively the first barrier that arboviruses affront to complete the transmission cycle, we decided to address the question of how this epithelium replenish its cells during the different challenges of blood feeding and infection. Previous descriptive reports of epithelial cell structure, function and midgut remodeling during metamorphosis [15,16,36] have shed some light on this process in mosquitoes, suggesting that the cell types described in other organisms, such as *Drosophila*, are also found in *A. aegypti*. Amongst the fully differentiated cells, the enterocytes were clearly distinguishable by their large nuclei size, abundance and localization. However, due to the current lack of mosquito specific markers for other differentiated and progenitor cells, like ee’s and EB’s, these cells were not properly identified in mosquitoes. Nonetheless, ISC hallmark capacity is to undergo mitosis, which can be marked using antibodies for phosphorylated histone 3. This allowed us to successfully identify the presence of ISC in the epithelium, and to quantify the number or cells dividing in the different conditions evaluated **(Fig 1A-B)**.

In the life history of mosquitoes, blood feeding represents a dramatic change from a sugar diet to ingestion of a large protein-rich meal. This transition imposes challenges to midgut homeostasis that are not faced by non-hematophagous insects. Knowledge about the mechanisms involved in the maintenance of midgut cellular integrity and homeostasis upon blood feeding or stress conditions is limited not only for *A. aegypti*, but also for other important vectors. In this study, we show unique properties of the mosquito midgut, suggesting that the regulation of epithelial cell proliferation is tightly regulated to allow proper handling of both chemical and biological sources of stress, including DENV infection, that occur during and after blood digestion. Based on these findings, we suggest that this regulation of midgut homeostasis is an important determinant of viral infection dynamics in the vector gut.

The maximal digestion rate is attained 24 hours after a blood meal [37]. Despite the dramatic increase of the microbiota, approximately 1000 times the levels before a meal [5], mosquitoes seem to maintain midgut epithelial cell turnover controlled **(Fig 1C and D)**. One explanation for this is the physical separation between the bolus and the epithelium by the PM, as in *Drosophila*. The PM is a thick extracellular layer composed mostly of chitin fibrils and glycoproteins that is gradually formed after a blood meal and surrounds the blood bolus, creating a physical separation from the midgut epithelium [7,8]. To preserve homeostasis, the PM establishes a selective barrier, permeable to nutrients and digestive enzymes but acting as a first line of defense against harmful agents. We show here that when the midgut epithelium was exposed to pathogenic bacteria ingested with the blood meal, thus before PM formation, there was a marked increase of mitosis (**Fig 2B**). More importantly, inhibition of the PM formation also resulted in elevated mitotic cell counts (**Fig 2C**). Treating insects with antibiotics abolished the mitosis upregulation promoted by chitin synthesis inhibition, further demonstrating that the contact of the blood bolus itself was not the determining factor to the increase mitotic cell numbers, but instead, the consequent exposure of the gut epithelium to the indigenous bacterial microbiota present in the lumen was the predominant event that elicited this response. In this way, the compartmentalization of the bolus may allow the enterocytes to minimize their exposure to deleterious agents, and it results in reduced need to shed and replenish damaged cells.

ROS production by midgut cells represents a major innate immunity effector mechanism that is involved in the control of the microbiota. However, ROS can also damage host cells, and thus, a proper balance between ROS production and microbial suppression is essential for the health of the host itself [31–34,38]. Here, we show that production of ROS was activated when PM formation was blocked and that this effect can be prevented by antibiotics (**Fig 2D**). Therefore, we propose that the signaling mechanism that leads to increased mitosis after exposure to indigenous bacteria is the production of ROS by the intestinal cells, as a defensive, yet possibly damaging, response (**Fig 2**).

The midgut epithelial cells are the first to support viral replication within the mosquito vector and several studies have addressed the immune response of the mosquito to the virus. Additionally, it is well-established that changes in ROS production in the midgut impact not only innate immunity responses against bacteria, but can also affect the mosquito ability to transmit human pathogens [5,39–42]. Despite this comprehensive knowledge about infection-related processes that occur within midgut cells, little is known about the cell turnover prior to and after infection. It was our intention to evaluate if this natural process of the midgut epithelium was different between mosquito strains with different degrees of susceptibility to DENV. Rockefeller (Rock) and Orlando (Orl) strains are susceptible and refractory strains respectively; however, under normal (sugar fed) conditions, they possess similar levels of mitotic cells (**S2 Fig**) Interestingly, the Orl strain possesses higher levels of mitosis than the Rock strain 24 hours after the blood meal (**Fig 3A-B**). This increased number of mitotic cells, is restricted to this specific time window, as 48 hours after the feeding, the numbers are no longer significantly different. This fact becomes relevant when the timeline is superposed to the timeline of the initial viral infection [43]. This becomes more apparent, when the numbers of mitotic cells on the susceptible Rock strain increase after 5 days, in a consistent timeline to the establishment of a successful infection with higher levels of infected cells, which is not observed in Orl strain that constrains the infection. In day 7, when the viruses normally leave the midgut to infect other tissues [43], the mitotic rate is reduced to levels compared of non-infected sugar-fed midguts (Fig 3C). Transcriptomic analyses of mosquito strains with different degrees of susceptibility to DENV revealed that some genes associated with cellular proliferation, growth and death are differentially expressed in refractory strains, upon DENV infection [44–47]. However, this has not been directly associated to midgut regeneration in these studies. In addition, the increased expression and activation of a variety of apoptotic cascade components in the midgut after viral infections implicate apoptosis as part of the *A. aegypti* defense against arboviruses [24,25,27]. Altogether, these studies pointed to the significant importance of cell replenishing in the midgut epithelium to vector competence. Because of that, we decided to target the Notch pathway through RNAi; to disturb the normal regenerative process of the epithelium. Amongst the proteins involved in this pathway, the ligand Delta was an excellent candidate for RNAi because it is upstream of the Notch signaling pathway and is considered a marker of ISC [19]. Induction of RNAi by injection of dsDelta in adult females, resulted in the silencing of the Notch ligand Delta, interrupted the normal cycle of cell replenishment resulted on reduced cell division (**Fig 4B and C**), as previously reported by Guo and Ohlstein (2015) in *Drosophila* and by VanDussen et al, 2012, in mice. As knockdown of Delta resulted on increased DENV2 viral titers in refractory strain **(Fig 4D)**, this suggested that cell regeneration is also a contributing factor to the modulation of viral infection and consequently to refractoriness. In addition to this result, we pre-treated mosquitoes of the susceptible strain (Rockefeller) with DSS, to induce cell division. Likewise, we found that the increase in mitosis was able to expand refractoriness of these mosquitoes. Our data shows for the first time that the ability to replenish the epithelial differentiated cells, by ISC engagement in tissue regeneration, is an important aspect of the mosquito’s antiviral response in these strains. Furthermore, these results revealed that the involvement of the Notch signaling pathway in midgut cell proliferation is also conserved in *A. aegypti*. Additional work is required to detail the mechanism by which Delta-Notch signaling interferes in midgut cell proliferation in the midgut of *A. aegypti*. The role of other pathways previously shown to regulate progenitor cell proliferation and differentiation in *Drosophila* and mammalians, such as the Hippo, JAK-STAT and other pathways, may also reveal key connections between intestinal cell replenishment and vectorial competence.

The first 24–48 h after ingestion of virus infected blood are considered the most critical for determining vector competence of a given mosquito (reviewed in [48]). Accordingly, we propose that the mitotic events in the early stages of infection (e.g., 24 h after viral ingestion) occur when the number of infected cells is still low and the capacity to eliminate damaged cells prevents viral spreading, and therefore must be effective to limit the infection. The number of mitotic cells of the refractory strain midgut at this initial time point is higher than in the susceptible strain, implicating this as a likely determinant for refractoriness **(Fig 4A and B)**. The differences observed in the total number of mitotic cells and in the pattern of recovery between Rockefeller and Orlando strains may suggest more extensive damage in the midgut of the susceptible mosquitoes caused by virus infection. However, the correlation between viral infection progression, cell damage and regenerative responses in the early infection remains to be investigated.

In conclusion, our data suggest that the midgut infection by DENV is favored by delayed midgut renewal in a permissive mosquito strain and that refractoriness would be supported, at least partly, by the capacity to promptly activate the ISC division program. At the present time, Dengue, Chikungunya and Zika viruses are widespread across the globe, and the understanding of the multiple factors affecting virus infection within the mosquito is crucial. The fact that faster cell renewal could be related to refractoriness adds up a new factor to be considered among the many determinants of vector competence and opens up the spectrum of the vector physiological events that are important when studying viral transmission. Future research is required to test if other DENV refractory field strains also possess differential tissue homeostatic properties and if a similar mechanism will occur in other arboviral infections. These findings reveal a new path towards a better understanding of vector competence, and may support the development of alternative strategies of virus transmission control.

Finally, these results highlight that the rate of midgut cell renewal should be taken into account when choosing mosquito strains for vector control strategies that use population replacement, such as SIT or *Wolbachia* based methodologies.

## Materials and Methods

### Ethics statement

All experimental protocols and animal care were carried out in accordance to the institutional care and use committee (Comitê para Experimentação e Uso de Animais da Universidade Federal do Rio de Janeiro/CEUA-UFRJ) and the NIH Guide for the Care and Use of Laboratory Animals (ISBN 0–309-05377-3). The protocols were approved under the registry CEUA-UFRJ #155/13. All animal work at JHU was conducted in strict accordance with the recommendations in the Guide for the Care and Use of Laboratory Animals of the National Institutes of Health (NIH), USA. The protocols and procedures used in this study were approved by the Animal Care and Use Committee of the Johns Hopkins University (Permit Number: M006H300) and the Johns Hopkins School of Public Health Ethics Committee.

### Rearing of *A. aegypti* mosquitoes

The *Aedes aegypti* (Red Eye strain) were raised at the insectary of UFRJ under a 12-hour light/dark cycle at 28°C and 70–80% relative humidity. The adults were maintained in a cage and given a solution of 10% sucrose *ad libitum* unless specified otherwise. The *Aedes aegypti* (Rockefeller and Orlando strains) were raised at the insectary of JHU under a 12-hourlight/dark cycle, at 27°C and 95% humidity. The adults were maintained in a cage and given a solution of 10% sucrose *ad libitum*. The adult females were dissected at different times after blood feeding for the experiments.

### Mosquito antibiotic treatment

The mosquitoes were rendered free of cultivable bacteria by maintaining them on a 10% sucrose solution with penicillin (100 μ/mL), and, streptomycin (100 μg/mL) from the first day post-eclosion until the time of dissection post blood feeding.

### Mosquito meals

The *Aedes aegypti* mosquitoes from the Red Eye strain (four- to seven-days-old) were artificially fed with heparinized rabbit blood. The feeding was performed using water-jacketed artificial feeders maintained at 37°C and sealed with parafilm membranes. The insects were starved for 4-8 hours prior to the feeding. Unfed mosquitoes were removed from the cages in all the experiments.

The oxidative challenge was provided by addition of 500 μM of paraquat (ChemService, West Chester, PA, USA) to the blood meal. As an antioxidant treatment, 50mM of ascorbic acid (neutralized to pH 7.0 with NaOH) was also added to blood. The mosquitoes were orally infected by *Serratia marcescens* BS 303 strain or *Pseudomonas entomophila* L48 strain at a concentration of 10^5^ bacteria/mL of blood. Briefly, overnight cultures were used either live or after heat inactivation. Inactivation of *P. entomophila* was done by heating at 98°C for 1 hour. Live and heat-killed bacteria were all pelleted after OD600 measurements to achieve final concentration of 10^5^ bacteria/mL of blood. The media supernatant was discarded and the cell pellet was resuspended in blood previous to the mosquito feeding. The compound diflubenzuron (DFB) (0.4 g/L), a well-known chitin synthesis inhibitor, was added to the blood meal to prevent the peritrophic matrix establishment [30].

To stimulate ISC proliferation and midgut regeneration [18], the mosquitoes were fed with 1% DSS (dextran sulfate sodium salt 6.5-10 kDa, Sigma, St. Louis, MO, USA) dissolved in 10% sucrose for 2 days before infection. Twelve hours prior to infection, the DSS-sucrose solution was substituted with a 10% sucrose solution to remove residual DSS from the midgut content. The control mosquitoes were fed with 10% sucrose only. The infection with DENV was carried out as described in the following sections.

### Proliferation and mitotic cells quantification

The quantification of mitosis in whole midgut tissues was performed by PH3 labeling as described elsewhere [49]. Briefly, female adult mosquitoes were dissected in PBS. Midguts were fixed in PBS with 4% paraformaldehyde for 30 minutes at room temperature. Samples were washed in PBS for 2 times of 10 minutes each. Then the tissues were permeabilized in PBS with 0.1% X-100 (for 15 min at room temperature) and blocked in a blocking solution containing PBS, 0.1% Tween 20, 2.5% BSA and 10% normal goat serum for at least 30 min at room temperature. All samples were incubated with primary antibody mouse anti-PH3 (1:500, Merck Millipore, Darmstadt, Germany). After washing 3 times of 20 minutes each in washing solution (PBS, 0.1% Tween 20, 0.25% BSA), samples were incubated with secondary goat anti-mouse antibody conjugated with Alexa Fluor 488 or 546 (Thermo Fisher Scientific, MA, USA) for at least 1 hour at room temperature at a dilution of 1:2000. DNA was visualized with DAPI (1mg/ml, Sigma), diluted 1:1000. The gut images were acquired in a Zeiss Observer Z1 with a Zeiss Axio Cam MrM Zeiss, and the data were analyzed using the AxioVision version 4.8 software (Carl Zeiss AG, Germany). Representative images were acquired using a Leica SP5 confocal laser-scanning inverted microscope with a 20X objective lens. Images were processes using Las X software.

### WGA and Phalloidin staining

Midguts from insects that were fed on naive blood or blood with DFB were dissected 24 h after feeding and fixed in 4% paraformaldehyde for 3 h. All of the midguts were kept on PBS-15% of sucrose for 12 h and then in 30% sucrose for 30 h. After a 24-h infiltration in OCT, serial microtome 14-lm-thick transverse sections were obtained and collected on slides that were subsequently labeled with the lectin WGA (Wheat Germ Agglutinin; a lectin that is highly specific for N-acetylglucosamine polymers) coupled to fluorescein isothiocyanate (FITC). The slides were washed 3 times in PBS buffer containing 2 mg/mL BSA (PBSB). The samples were then incubated in 50mM NH4Cl/PBS for 30 min; in 3% BSA, 0.3% Triton X-100 PBS for 1 h; and in PBSB solution with 100 mg/mL WGA-FITC (EY Laboratories) for 40 min. The slides were then washed three times with PBSB and mounted with Vectrashield with DAPI mounting medium (Vector laboratories). The sections were acquired in an Olympus IX81 microscope and a CellR MT20E Imaging Station equipped with an IX2-UCB controller and an ORCAR2 C10600 CCD camera (Hammamatsu). Image processing was performed with the Xcellence RT version 1.2 Software.

Midguts from insects that were fed on blood alone or blood with DENV-2 were dissected 5 days after feeding and fixed in 4% paraformaldehyde using the same protocol as for mitotic cell quantification. After the secondary antibody incubation washes, 30 min incubation with phalloidin 1:100 (1uL) in 98uL blocking solution, along with the DAPI (1:100) was done at room temperature protected from light. Samples were washed twice, for 5 minutes (stationary, room temperature, protected from light) in 0.5mL washing solution and then onto slides with VectaShield. Images (z-stack of 0.7 μm slides) were taken on a Zeiss LSM700 laser scanning confocal microscope at the Department of Cell Biology at JHU with a 20X objective lens and processed using Zeiss Zen Black Edition software.

### ROS detection in the midgut

The mosquito midguts were dissected in PBS 24h after feeding and incubated with 50μM of dihydroethidium (hydroethidine; DHE; Invitrogen) diluted in Leibovitz-15 media supplemented with 5% fetal bovine serum for 20 min at room temperature in the dark. The incubation media was gently removed and replaced with a fresh dye-free media. The midguts were positioned on a glass slide, and the oxidized DHE-fluorescence was observed by a Zeiss Observer Z1 with a Zeiss Axio Cam MrM Zeiss using a Zeiss-15 filter set (excitation BP 546/12; beam splitter FT 580; emission LP 590) (Carl Zeiss AG, Germany) [5,50].

### RNA extraction and qPCR analysis

For the qPCR assays, the RNA was extracted from the midgut using TRIzol (Invitrogen, CA, USA) according to the manufacturer’s protocol. The complementary DNA was synthesized using the High-Capacity cDNA Reverse transcription kit (Applied Biosystems, CA, USA). The qPCR was performed with the StepOnePlus Real Time PCR System (Applied Biosystems, CA, USA) using the Power SYBR-green PCR master MIX (Applied Biosystems, CA, USA). The Comparative Ct method [51,52] was used to compare the changes in the gene expression levels. The *A. aegypti* ribosomal S7 gene was used as an endogenous control [53]. The oligonucleotide sequences used in the qPCR assays were S7 (AAEL009496-RA): S7_F: GGGACAAATCGGCCAGGCTATC and S7_R: TCGTGGACGCTTCTGCTTGTTG; Delta (AAEL011396), Delta_Fwd: AAGGCAACTGTATCGGAGCG and Delta_Rev: TATGACATCGCCAAACGTGC.

### Gene silencing

Two- to three-day old mosquito females (Rockefeller and Orlando) were cold anesthetized and 69 nL of 3 μg/μL dsRNA solution was injected into the thorax. Three days after injection, the mosquitoes were infected with DENV. Mosquito midguts were collected after 24h for real time PCR and after 5 days for mitosis assay or DENV infection analysis. The HiScribe T7 *in vitro* transcription kit (New England Biolabs) was used to synthesize the dsRNA. The unrelated dsGFP was used as a control, and the silencing efficiency was confirmed through qPCR. To generate dsDelta, the following oligonucleotides (containing the T7 polymerase-binding site) were used:

dsDelta_Fwd: <u>GTAATACGACTCACTATAGGG</u>AGCAAGCCTAACGAGTGCAT
dsDelta_Rev: <u>GTAATACGACTCACTATAGGG</u>TTCCTTCTCACAGTGCGTCC

### Dengue virus propagation and mosquito infections

The DENV-2 (New Guinea C strain) was propagated for 6 days in C6/36 cells maintained in complete MEM media supplemented with 10% fetal bovine serum, 1% penicillin/streptomycin, 1% non-essential amino acids and 1% L-glutamine. The virus titer was determined by plaque assay as 10^7^ PFU/mL [54]. The females were infected through a blood meal containing: one volume of virus, one volume of human red blood cells (commercial human blood was centrifuged and the plasma removed), 10% human serum and 10% 10 mM ATP. Unfed mosquitoes were removed from the cages. The midguts were dissected at 5 days post-blood meal and stored individually in DMEM at −80°C until used.

For DENV-4 (Boa Vista 1981 strain) propagation, the virus was cultivated 6 days in C6/36 cells maintained in Leibovitz-15 media supplemented with 5% fetal bovine serum, 1% non-essential amino acids,1% penicillin/streptomycin and triptose (2.9 g/L) [55]. The virus titer was determined by plaque assay as 10^7^ PFU/mL. The females that were pre-treated with DSS or regular sucrose (control) were infected using one volume of rabbit red blood cells and one volume of DENV-4. The midguts were dissected at 7 days after infection and stored individually in DMEM at −80°C until used.

### Plaque assay

The plaque assay was performed as previously described [28]. The BHK-21 cells were cultured in complete DMEM media, supplemented with 10% fetal bovine serum, 1% penicillin/streptomycin and 1% L-glutamine. One day before the assay, the cells were plated into 24 wells plates at 70-80% confluence. The midguts were homogenized using a homogenizer (Bullet Blender, Next, Advance) with 0.5mm glass beads. Serial dilutions (10 folds) were performed, and each one was inoculated in a single well. The plates were gently rocked for 15 min at RT and then incubated for 45 min at 37°C and 5% CO_2_. Finally, an overlay of DMEM containing 0.8% methylcellulose and 2% FBS, was added in each well, and the plates were incubated for 5 days. To fix and stain the plates, the culture media was discarded and a solution of 1:1 (v:v) methanol and acetone and 1% crystal violet was used. The plaque-forming units (PFU) was counted and corrected by the dilution factor.

### Statistical analysis

Unpaired Student’s t-tests were applied where comparisons were made between two treatments or two different mosquito strains, as indicated in the figure legends. Mann-Whitney U-tests were used for infection intensity and chi-square tests were performed to determine the significance of infection prevalence analysis. All statistical analyses were performed using GraphPad 5 Prism Software (La Jolla, United States).

## Acknowledgments

We thank all members of the Laboratory of Biochemistry of Hematophagous Arthropods, especially Jaciara Loredo, Mônica Sales and S.R. Cassia for providing technical assistance. We also thank Dr. Helena Araujo (ICB, UFRJ) for all the technical advice and assistance with the microscopy experiments. We would like to thank the Johns Hopkins Malaria Research Institute Insectary.

